# Integrated Systems-Analysis of the Human and Murine Pancreatic Cancer Glycomes Reveal a Tumor Promoting Role for ST6GAL1

**DOI:** 10.1101/2021.03.10.434864

**Authors:** Emma Kurz, Shuhui Chen, Emily Vucic, Gillian Baptiste, Cynthia Loomis, Praveen Agarwal, Cristina Hajdu, Dafna Bar-Sagi, Lara K. Mahal

## Abstract

Pancreatic ductal adenocarcinoma (PDA) is the 3^rd^ leading cause of cancer-death in the U.S.. Glycans, such as CA-19-9, are biomarkers of PDA and are emerging as important modulators of cancer phenotypes. Herein, we utilized a systems-based approach integrating glycomic analysis of human PDA and the well-established KC mouse model, with transcriptomic data to identify and probe the functional significance of aberrant glycosylation in pancreatic cancer. We observed both common and distinct patterns of glycosylation in pancreatic cancer across species. Common alterations included increased levels of α-2,3- and α-2,6-sialic acids, bisecting GlcNAc and poly-LacNAc. However, core fucose, which was increased in human PDAC, was not seen in the mouse, indicating that not all human glycomic changes can be modeled in the KC mouse. In silico analysis of bulk and single cell sequencing data identified ST6GAL1, which underlies α-2,6-sialic acid, as overexpressed in human PDA, concordant with histological data. Enzymes levels correlated with the stage of clinical disease. To test whether ST6GAL1 promotes pancreatic cancer we created a novel mouse in which a pancreas-specific genetic deletion of this enzyme overlays the KC mouse model. Analysis of our new model showed delayed cancer formation and a significant reduction in fibrosis. Our results highlight the importance of a strategic systems-approach to identifying glycans whose functions can be modeled in mouse, a crucial step in the development of therapeutics targeting glycosylation in pancreatic cancer.

**SIGNIFICANCE:** Pancreatic ductal adenocarcinoma (PDA) is the 3^rd^ leading cause of cancer-death in the U.S.. Glycosylation is emerging as an important modulator of cancer phenotype. Herein we use a systems-approach integrating glycomics of human PDA and a well-established PDA mouse model with transcriptomic data to identify ST6GAL1, the enzyme underlying α-2,6-sialic acid, as a potential cancer promoter. A pancreatic specific ST6GAL1 knockout in the KC mouse showed delayed cancer formation and a reduction in fibrosis. Our results highlight the importance of a strategic systems-approach to identifying glycans whose functions can be modeled in mouse, a crucial step in the development of therapeutics targeting glycosylation in pancreatic cancer.

## INTRODUCTION

The survival rate for pancreatic ductal adenocarcinoma (PDA) beyond 5 years is very low. This disease is the 3^rd^ leading cause of cancer-related death in the U.S., with few treatment options once malignant transformation has occurred. Growing evidence has identified altered glycosylation as a hallmark of solid tumor cancers (1). Glycosylation contributes to multiple facets of both cancer initiation (e.g., sustained proliferative signaling, resistance to cell death, etc.) and progression (e.g., invasion, metastasis, tumor-promoting inflammation, etc.) (1). Several studies have identified the glycan sialyl Lewis A (sLe^A^), also known as carbohydrate antigen CA 19-9, as a biomarker of pancreatic cancer and as a prognostic indicator (2, 3). Heterologous expression of human enzymes that can biosynthesize CA-19-9 in a mouse model increased pancreatic inflammation, pointing to a potential role of this epitope in disease initiation (4). Pancreatic cell culture models have also shown a potential role for glycosylation in cell stemness, invasiveness and drug resistance (5–8).

Much of our knowledge of cancer biology comes from mouse models. Glycosylation has both conserved and species-specific components (9). Many glycans and glycosylation enzymes are conserved across species, but can have distinct biological functions in different organisms. For example, mutations and deletions in ST3GAL5, an enzyme that biosynthesizes GM3, are responsible for a spectrum of severe epileptic disorders in humans (10). However, a mouse model in which ST3GAL5 (also known as GM3 synthase) is knocked out does not show this phenotype (11, 12). The lack of direct conservation between the mouse and human biology when centered on the glycome is a crucial issue in modeling this axis of human disease.

Herein, we use a systems-based approach integrating glycomic analysis of both human PDA and a mouse model with transcriptomic data to identify and probe the functional significance of aberrant glycosylation in pancreatic cancer. Specifically, we show that α-2,3- and α-2,6-sialic acids, bisecting GlcNAc and poly-LacNAc are significantly upregulated in both human PDA and the KC mouse model, while other glycans are only seen in one of the two sample sets. This mouse model, in which the expression of mutant KRas observed in pancreatic cancer patients (KRAS^G12D^) is driven by the pancreatic lineage specific p48^Cre^, is considered an accurate representation of the axis of human disease. Using transcriptomic data we identified specific glycosyltransferases driving these glycopatterns, narrowing our focus upon ST6GAL1, the main enzyme underlying α-2,6-sialic acid. In-silico analysis of single cell sequencing data pointed to an increase of ST6GAL1 in the cancerous ductal compartment. Patients with higher levels of ST6GAL1 transcripts had lower overall survival. Multiplex immunofluorescence confirmed that ST6GAL1 protein is significantly upregulated in human disease and correlates with advanced tumor stage. To test whether this enzyme is a promoter of pancreatic cancer, we created a pancreas-specific genetic deletion of ST6GAL1 in the KC mouse model (ST6KC). Analysis of this model showed delayed cancer formation and a significant reduction in fibrosis. Our results highlight the importance of a strategic systems-approach to identifying glycans whose functions can be modeled in mouse, a crucial step in the development of therapeutics targeting glycosylation in pancreatic cancer.

## RESULTS

### Human Pancreatic Cancer Displays Strong Shifts in the Glycome

To study whether pancreatic cancer displays an altered glycome from the normal pancreas, we utilized our dual-color lectin microarray technology (**Fig. 1a**, ***SI Appendix*, Fig. S1a**) (13, 14). Lectin microarrays employ carbohydrate-binding proteins with well-defined specificities to detect glycan changes between samples and have been used for cancer glycomics (15–18). For our analysis, we obtained 12 biopsy samples from non-cancerous pancreata and 12 from pancreatic adenocarcinoma (PDA) of patients (***SI Appendix,* Table S1**). Previous glycomic analysis on pancreatic cancer used “normal adjacent” tissue, but this is not considered representative of the non-transformed pancreas (19, 20). In brief, formalin-fixed paraffin embedded (FFPE) pancreatic tissue samples were solubilized and labeled with Alexa Fluor 555-NHS as previously described (***SI Appendix,* Table S2**) (15). Equal amounts of sample and orthogonally labeled reference (Alexa Fluor 647-labeled, pooled from all samples) were analyzed. The heatmap of lectins showing significant differences in binding between pancreatic cancer and normal tissue is shown in **Fig. 1b**, ***SI Appendix*, Fig. S1b, c**.

**Figure 1.**
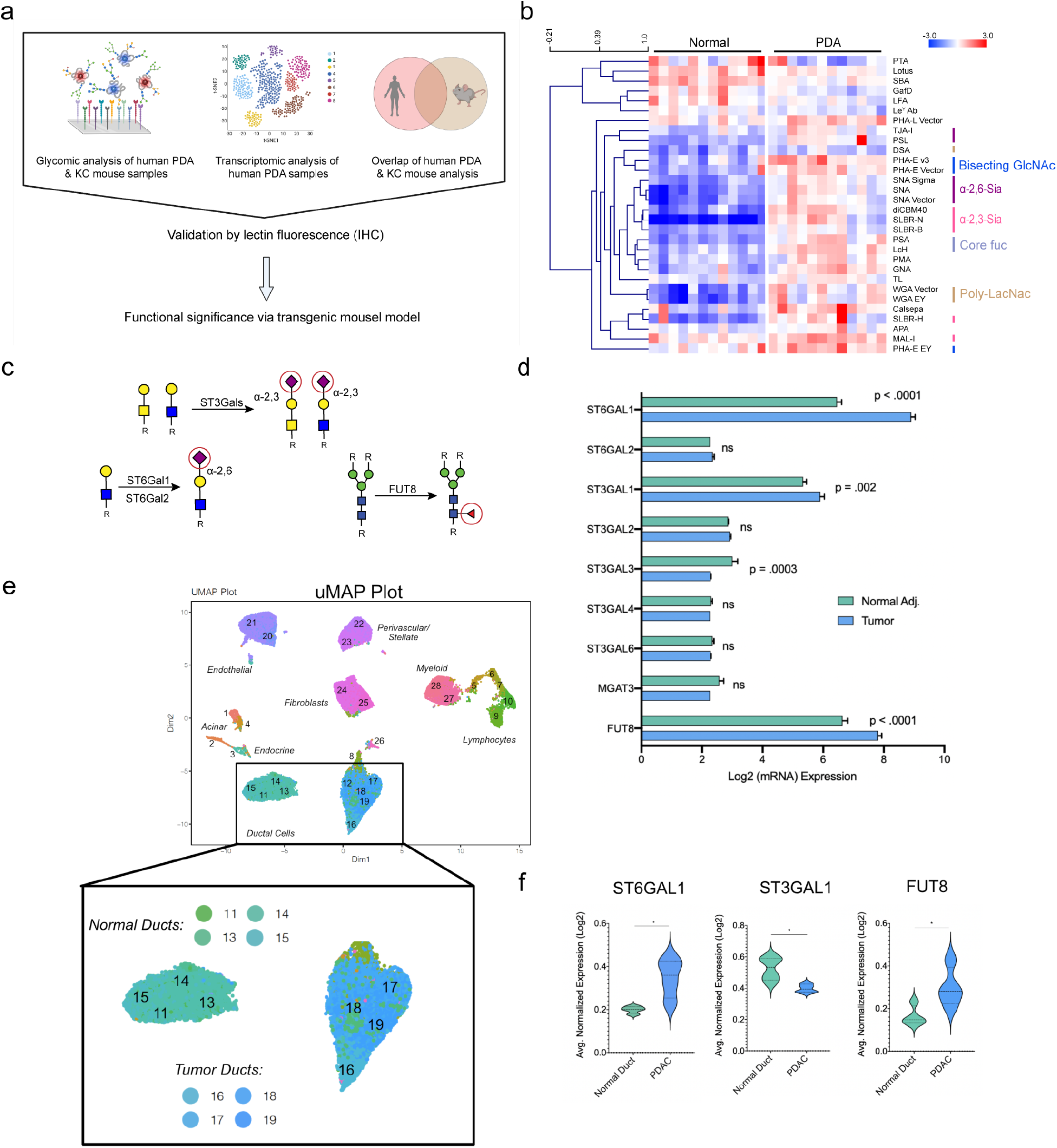
Integrated systems analysis points to ST6GAL1 and FUT8 as potential drivers of PDA. a) Schematic representing systems approach analysis of lectin protein and transcriptome analysis in mouse and human pancreatic cancer. b) Heatmap of human lectin microarray data present with significant lectins (*p*<0.05). Median normalized log_2_ ratios (Sample (S)/Reference(R)) were ordered by sample type (Normal, n = 12; PDA, n = 12). Red, log_2_(S) > log_2_(R); blue, log_2_(R) > log_2_(S). Lectins binding 〈-2,3-sialosides (pink), 〈-2,6-sialosides (purple), bisecting GlcNAc (navy), poly-*N*-Acetyl-D-Lactosamine (poly-LacNAc, brown), core fucose (slategrey) and Lewis antigens (turquoise) are highlighted to the right of the heatmap. c) Biosynthetic pathways for glycans underlying lectin signature are shown. Glycans are annotated following the Symbolic Nomenclature for Glycans (1). d) Transcriptomic analysis assessing the mRNA levels of select glycosyltransferases between normal adjacent and matched cancerous tissues within the same patient. e) uMAP plots representing cells isolated from PDAC (n=24) patients and normal pancreata (n=11) pooled on single cell-sequencing. Clusters representing normal ductal cells and tumor ductal cells are highlighted. f) Violin plot showing comparison of ST6Gal1 levels in Normal (green) vs. PDAC (blue) ductal clusters. Clusters in each group are combined. Violin plots showing comparison of ST6Gal1, FUT8, and ST3GAL1 levels in Normal (green) vs. PDAC (blue) ductal clusters. Clusters in each group are combined (ns: p>0.05; *: p<0.05)

We observed strong differences between the non-cancerous and cancerous pancreatic glycome, with greater than 1/3 of all lectins on the array showing statistically significant differences in binding. In PDA we observed significant increases in sialic acids, with changes in both α-2,6- (lectins: SNA, TJA-I, PSL, average increase: ~ 2.7 fold) and α-2,3-sialosides (diCBM40, SLBR-H, SLBR-B, SLBR-N, MAL-I, MAA, ~ 3.4 fold). Core fucosylation (PSA, LcH, ~ 2.7 fold), bisecting GlcNAc (PHA-E, ~ 2.2 fold) and polyLacNAc (WGA, DSA, LEA, ~ 2.1 fold) also increased relative to normal. This dramatic shift in the glycome in transformed pancreas strongly suggests a potential role for glycosylation in the etiology of this disease.

### Bulk Transcriptomic Analysis and Single Cell Sequencing Reveal ST6GAL1 and FUT8 are Enriched in Cancerous Ducts

We next sought to determine glycosyltransferases that underlie the observed glycomic signatures. Using publicly available bulk RNA sequencing (bulk-seq) data from three separate studies performed on the same platform, we interrogated the expression levels of select glycan biosynthetic enzymes (ST6GAL1, ST6GAL2 [α-2,6-sialylation], ST3GAL1, ST3GAL2, ST3GAL3, ST3GAL4, ST3GAL6 [α-2,3-sialyation], FUT8 [core fucose] and MGAT3 [bisecting GlcNAc], **Fig. 1c and d**). Comparison of human pancreatic cancer to adjacent normal pancreas showed statistically significant increases for ST6GAL1, ST3GAL1 and FUT8 in PDA. The PDA tumor and its microenvironment is composed of many cell types including ductal cells, immune cells, fibroblasts, and endothelia. We analyzed a separate cohort of publicly available single cell RNA sequencing (sc-RNAseq) data using iCellR to study cell type specific expression of upregulated genes from bulk-seq (ST6GAL1, ST3GAL1, FUT8) (21–23). uMAP clusters were defined by transcriptomic signatures typical of intra-pancreatic cell types (**Fig. 1e**, ***SI Appendix,* Fig. S2**). PDAC originates from exocrine tissue specifically composed of acinar and ductal cells. Strong ST6GAL1 expression was observed in endothelial, immune cell and cancerous ductal clusters. We observed significant enrichment of this α-2,6 sialic acid enzyme in tumor ducts compared to the normal pancreas (**Fig. 1f**, ***SI Appendix,* Fig. S3a**). The α-2,3 sialyltransferase ST3GAL1 was also observed in many cell types, however it was not enriched in cancer-specific ductal cells (**Fig. 1f**, ***SI Appendix,* Fig. S3b).** The core fucosylation enzyme, FUT8, showed strong expression in fibroblasts, endothelial cells and cancerous ductal cells, with significant enrichment in cancerous ducts when compared to normal ducts (**Fig. 1f**, ***SI Appendix,* Fig. S3c**). Our data analysis suggests that α-2,6 sialic acid and core fucosylation may be important to the formation and/or progression of cancerous ducts.

### The KC Mouse Model Replicates Some, But Not All of the Human PDA Glycomic Signature

To assess the functional impact of glycosylation on pancreatic cancer development and progression, we turned to a mouse model. Glycosylation has both conserved and species-specific components (9). Many glycans and glycosylation enzymes are conserved across species, but can have distinct biological functions in different organisms. For example, mutations and deletions in ST3GAL5, an enzyme that biosynthesizes GM3, are responsible for a spectrum of severe epileptic disorders in humans (10). However, a mouse model in which ST3GAL5 (also known as GM3 synthase) is knocked out does not show this phenotype (11, 12). To determine whether we can model the impact of α-2,6 sialic acid, core fucosylation or other glycomic changes observed in human pancreatic cancer in mice, we first determined whether these glycans were altered in the KC mouse model. This model, which is considered an accurate representation of the axis of human disease, uses a mutant KRas, observed in 90% of pancreatic cancer patients, to drive disease.

To investigate whether the KC model accurately represents changes observed in the human PDAC glycome, we analyzed pancreata from genetically engineered **KC** mice bearing a p48-specific activating KRAS mutation (**KC**: p48^Cre^; LSL^KRASG12D^). These mice develop dysplasia, pan-IN lesions, fibrosis, and eventually advanced cancer of the pancreas (24). Pancreata were harvested from 14-week old female (n= 8) and male (n=5) KC mice, along with appropriately age and gender matched litter mate controls (**Normal**: female, n = 4; male, n= 5). By this time point, significant levels of advanced pan-IN lesions and displacement of normal acinar tissue are observed. Formalin-fixed paraffin embedded (FFPE) pancreatic tissue samples were processed and analyzed using our dual-color lectin microarray technology (15). Heatmaps of lectins showing significant differences in binding between pancreatic cancer and normal tissue, separated by sex, are shown in **Figure 2**, ***SI Appendix***, **Figure 4**.

**Figure 2.**
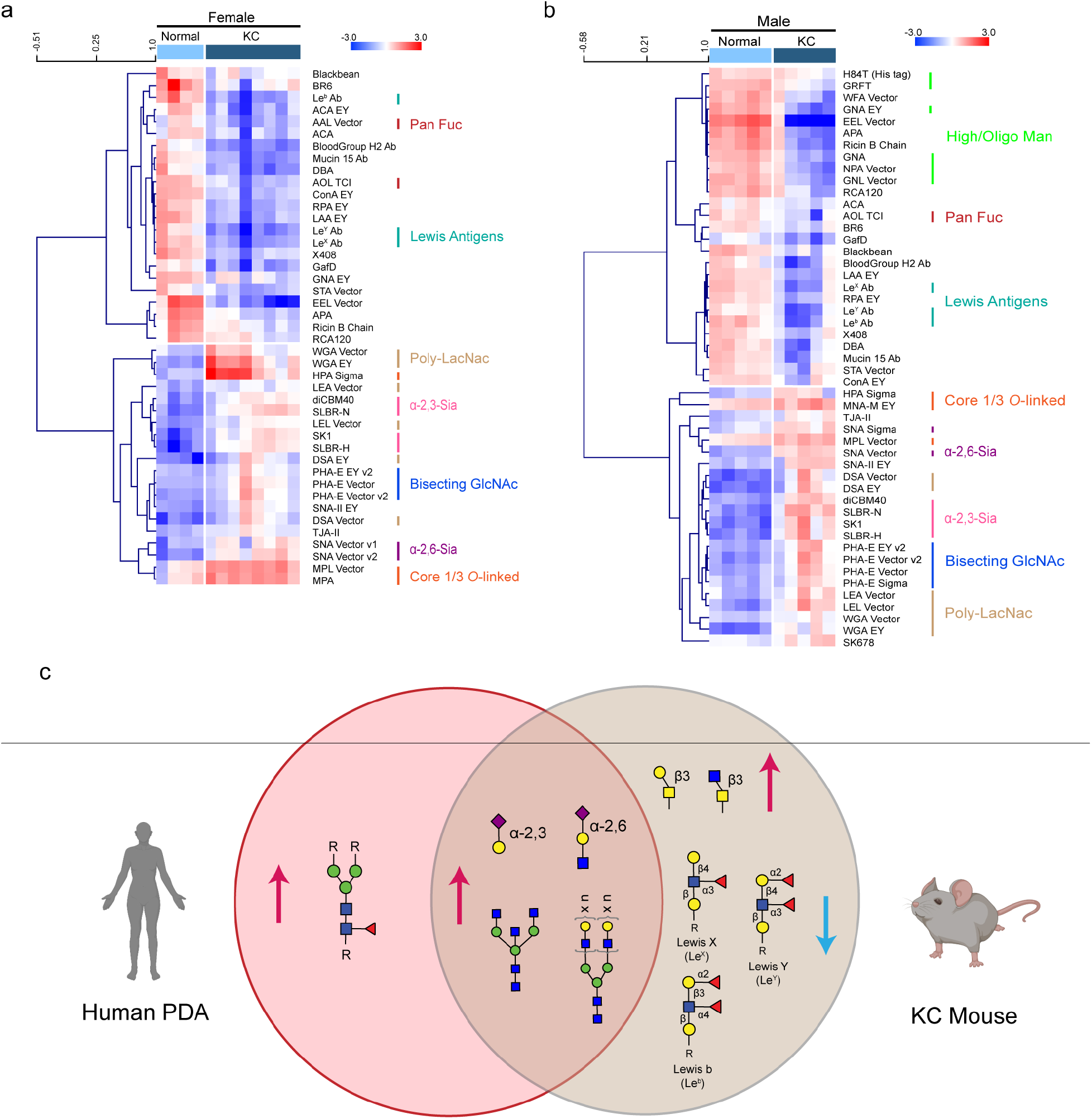
Glycomic analysis of KC mice shows both common and distinct glycopatterns in PDA. a) Heatmap of female mice lectin microarray data present with significant lectins (*p*<0.05). Median normalized log_2_ ratios (Sample (S)/Reference(R)) were ordered by sample type (Normal, n = 4; KC, n = 8). Red, log_2_(S) > log_2_(R); blue, log_2_(R) > log_2_(S). Lectins binding *α*-2,3-sialosides (pink), *α*-2,6-sialosides (purple), bisecting GlcNAc (navy), poly-*N*-Acetyl-D-Lactosamine (poly-LacNAc, brown), core 1/3 *O*-linked glycans (orange), pan fucose (red) and Lewis antigens (turquoise) are highlighted to the right of the heatmap. b) Heatmap of male mice lectin microarray data present with significant lectins (*p*<0.05). Median normalized log_2_ ratios (Sample (S)/Reference(R)) were ordered by sample type (Normal, n = 5; KC, n = 5). Red, log_2_(S) > log_2_(R); blue, log_2_(R) > log_2_(S). Lectins binding *α*-2,3-sialosides (pink), *α*-2,6-sialosides (purple), bisecting GlcNAc (navy), poly-*N*-Acetyl-D-Lactosamine (poly-LacNAc, brown), high/oligo-mannose (bright green), core 1/3 *O*-linked glycans (orange), pan fucose (red) and Lewis antigens (turquoise) are highlighted to the right of the heatmap. c) Summary of glycomic changes between the KC mouse model and human PDA. 〈-2,3- and 〈-2,6-sialic acid epitopes, bisecting GlcNAc and poly-LacNAc are conserved glycans. Glycans are drawn in the Symbolic Nomenclature for Glycomics (SNFG). Symbols are defined as follows: galactose (yellow circles), *N-* acetylgalactosamine (yellow squares), *N-*acetylglucosamine (blue squares), mannose (green circles) and sialic acid (purple diamonds).

Glycomic analysis shows gender-dependent and independent differences in pancreatic cancer in mouse. Focusing on common differences between sexes, we observe increases in sialic acids (2-4 fold) in the KC mouse pancreata. These changes were seen in α-2,6- (lectins: SNA) and α-2,3-sialosides (lectins: diCBM40, SLBR-H, SLBR-N, SK1). Similar dramatic increases were also observed in bisecting GlcNAc (PHA-E) and poly-*N*-Acetyl-D-Lactosamine (DSA, LEA, WGA). Additionally, we observed increases in core 1/3 *O*-linked glycans (MNA-M, MNA-G, MPA, AIA). In contrast, Lewis antigens, which are fucosylated epitopes, and overall fucose levels decrease in KC mice relative to normal controls (AOL, AAL, antibodies: Le^b^, Le^X^, Le^Y^). We see no evidence of an increase in core fucosylation (LcH, PSA, ***SI Appendix***, **Fig. 4**).

Only a subset of the overall glycan signature observed in the KC mouse model recapitulated the human PDAC data. (**Fig. 2c**). Conserved glycomic changes between the two species include α-2,6- and α-2,3-sialosides, bisecting GlcNAc and polyLacNAc. The core fucosylation signature, associated with FUT8, which is a strong signature in all of the human data, was not observed in the KC mouse model. Based on this and our previous transcriptomic analysis, we focused our attention on the α-2,6 sialylation signature and associated enzyme ST6GAL1.

### Levels of ST6Gal1 Correlate with Stage and Survival in Human Pancreatic Cancer

To look more deeply at α-2,6 sialylation in human pancreatic cancer we stained an orthogonal cohort of human PDA tumors and normal pancreata for ST6GAL1 and α-2,6- sialylation (SNA, **Fig. 3a,b**, ***SI Appendix*, Fig. S5a-b**). Consistent with both our lectin microarray data and transcriptomic analysis, we observed that α-2,6-sialic acid levels (as determined by SNA) and corresponding ST6GAL1 levels are both significantly upregulated in human cancer (**Fig. 3c,d**). Further analysis revealed that more advanced human pancreatic cancers (tumor-nodal-metastasis (TMN) Stage 3 or 4) display significantly higher levels of ST6GAL1 staining than Stage I, Stage II or normal controls (**Fig. 3e**). Although we observe an increase in SNA staining with early stages (i.e., Stage I and II), we do not see this increase in more advanced stages on our tissue microarray (***SI Appendix***, **Fig. 5c**). This may be due to the lower numbers of cases of Stage 3 and 4 cancer on the array. Upon closer examination of the stained tissues, we observed ST6GAL1 in both stromal infiltrates and transformed epithelial cells (***SI Appendix***, **Fig. 5d,e**). Epithelial cells encompass endocrine, acinar and ductal cells in the pancreas. Consistent with the sc-seq. analysis, epithelial expression of ST6GAL1 correlates more significantly with disease state than stromal expression (***SI Appendix***, **Fig. 5d,e**). High expression of ST6GAL1 is also significantly correlated with reduced overall survival in PDA in the TCGA dataset (**Fig. 3f**). This corroborates our findings that ST6GAL1 at the protein level is associated with higher disease stage.

**Figure 3.**
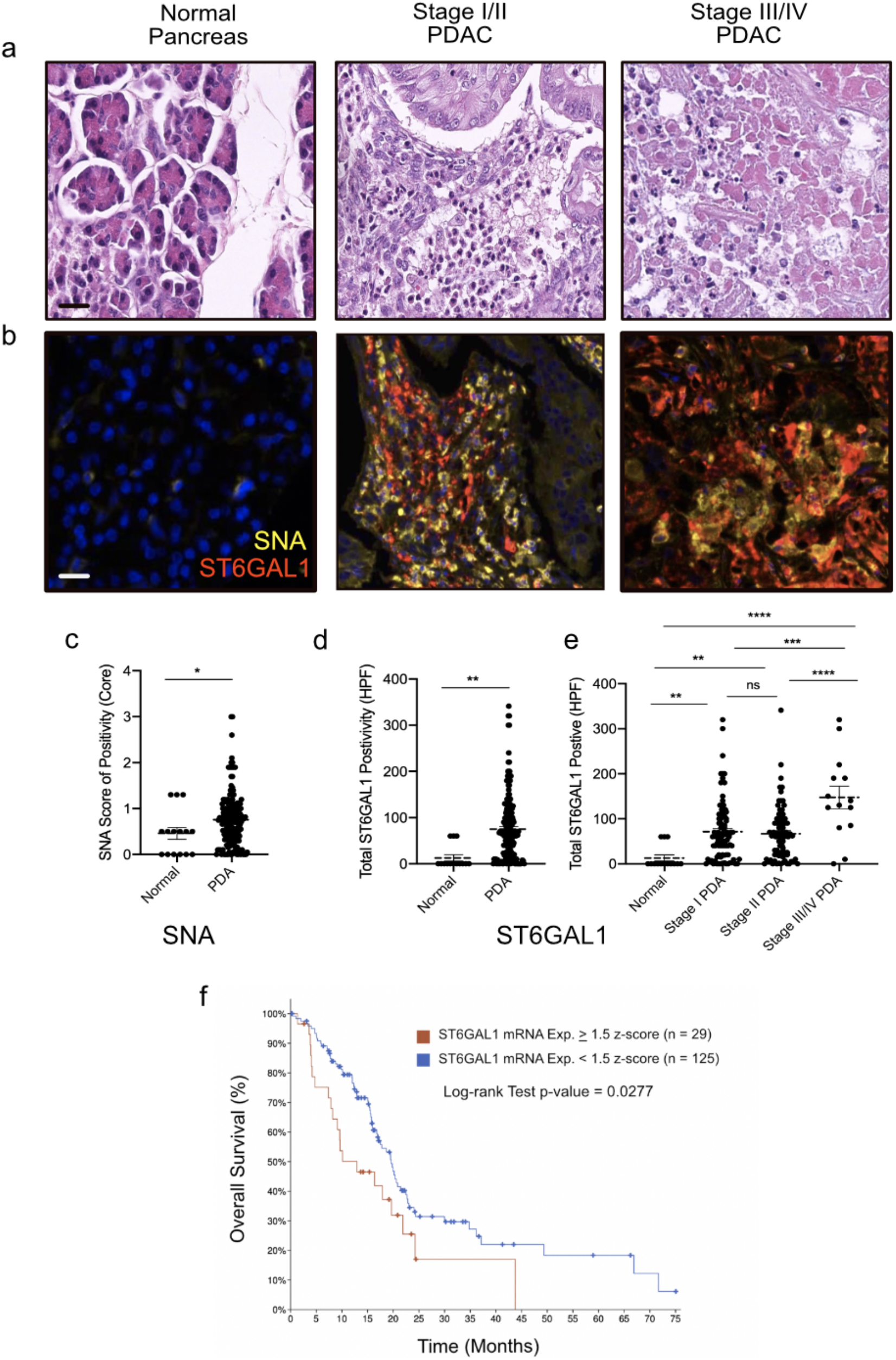
Profiling of ST6GAL1 and SNA in human pancreatic cancer shows association with stage and survival. a) H&E of normal pancreas (left), stage I pancreatic adenocarcinoma (center), and stage IV PDA (right) stained from a BioMAX human tissue microarray. b) Multiplex OPAL IF staining of SNA (yellow), ST6GAL1 (red) and DAPI (blue) on corresponding normal pancreas, Stage I, and Stage IV pancreatic adenocarcinoma from purchased BioMAX human tissue microarray. Scale bars represent 25 mm. c) Quantification of SNA positive cells per high powered field based on multiplex IF in normal versus all cancerous cases in human tissue microarray. d) Quantification of ST6GAL1 positive cells per high powered field in normal versus all cancerous cases in human tissue microarray. e) Distribution of the total ST6GAL1 positive cells per high power field across normal and each stage of pancreatic cancer based on multiplex IF imaging of human pancreatic cancer tissue microarray. f) Survival analysis of (n=154) verified human PDAC tumors from TCGA Pan-Cancer Clinical Data Resource (TCGA-CDR) processed in cBioPortal and separated by relative mRNA z-score for ST6GAL1 expression. (Log-rank Test p-value < 0.05 = statistical significance). (ns: p > 0 .05, *: p < 0.05, ** : p < 0.01, *** : p < 0.001, **** : p < 0.0001).

### Pancreas Specific Deletion of ST6GAL1 Slows Cancer Development and Progression

To determine whether ST6GAL1 plays a role in the development or progression of pancreatic cancer, we generated a novel ST6GAL1flx/flx KC mouse (ST6KC). In these mice, a pancreatic-lineage specific p48-Cre drives both the expression of mutant KRAS G12d and knockout of ST6GAL1 (**Fig. 4a**). To validate the ST6GAL1 knockout, we performed staining for ST6GAL1 in pancreata from the ST6KC mice at 14 weeks of age. As expected, we observed a significant loss of both ST6GAL1 protein and concomitant α-2,6-sialylation (as seen by SNA staining) in ductal and acinar cells (**Fig. 4b-c**). We next examined whether the knockout altered the histological profile of pancreata from ST6KC as compared to KC mice. We observed significant preservation of normal acinar area in the ST6KC compared to KC mice (**Fig. 4d**, *p* < 0.01). Examination of fibrosis, as quantified by Trichrome and Gomore staining showed a significant reduction in ST6KC mice (**Fig. 4e**, *p* < 0.01). Fibrosis is a hallmark of pancreatic cancer formation and severity. It is known to contribute to disease progression, immune suppression, and resistance to therapeutic interventions in both mouse and human disease (25). Overall, our model demonstrates that targeted deletion of ST6GAL1 has a clear protective effect against pancreatic cancer formation and progression.

**Figure 4.**
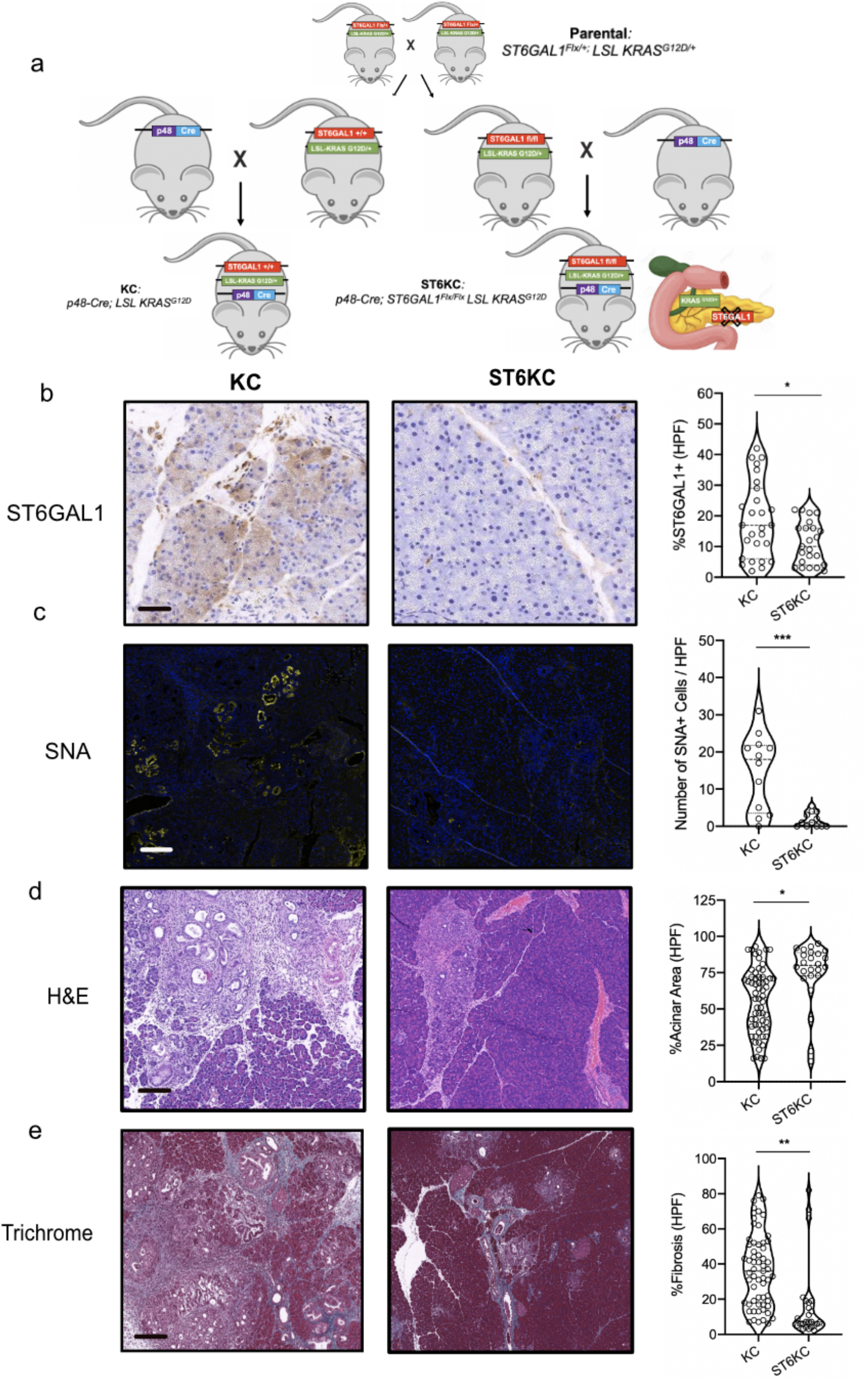
Pancreas-specific deletion of ST6GAL1 reduces disease burden in murine PDA. a) Breeding schematic illustrating the generation of novel ST6KC mice. Offspring of parental strains crossed into p48 Cre mice drive induction of mutant KRAS^G12D^ and deletion of ST6GAL1 under the same promoter. b) Immunohistochemistry staining for ST6GAL1 in 14 week littermate FFPE pancreata from KC and ST6KC mice (n=5 per group). Number of ST6GAL1 positive epithelial cells per high powered field is quantified on the right, each dot is a single field. Scale bar represents 80 mm. c) SNA staining of 14 week old FFPE pancreata from KC and ST6KC mice (n=3 per group). Scale bar represents 100 mm. d) H&E of 14 week old FFPE pancreata from KC and ST6KC mice (n=5 per group). Percent of preserved normal pancreas area is quantified per high power field on right Scale bar represents 200 mm. d) Trichrome and Gomore (blue) stain of FFPE pancreata from 14 week KC and ST6KC mice (n=5 per group). Percent of collagen deposition fibrosis is quantified per high power field on right. Scale bar represents 200 mm. (ns = p > 0.05, * = p < 0.05, ** = p < 0.01, *** = p <0 .001).

## CONCLUSIONS

Pancreatic cancer is one of the deadliest cancers and is predicted to be the 2^nd^ leading cause of cancer-related death by 2030 (26). While the incidence rate of pancreatic cancer has increased, the 5-year survival rate remains low, with few long-term survivors and high rates of therapeutic resistance (27). Long known as a hallmark of cancer, changes in glycosylation are emerging as crucial modulators of cancer phenotype. Herein, we applied a systems-based approach integrating lectin microarray analysis of human samples, mouse models and in silico analysis of both bulk and single cell sequencing data to investigate the role of glycosylation in promoting pancreatic cancer. Our approach guided our modeling of the impact of glycosylation on pancreatic cancer, enabling testing of ST6GAL1 in the KC mouse model and identifying it as a promoter of this disease.

Although glycomic analysis of human samples identified dramatic shifts in glycosylation in pancreatic cancer, only a select set of these were recapitulated in the KC mouse model. These include prominent increases in α-2,3- and α-2,6-sialosides, bisecting GlcNAc, and poly-LacNAc epitopes. Similar shifts in sialylation and bisecting GlcNAc levels were observed in work comparing the glycome of pancreatic cancer to patient-matched adjacent-tissue (19). However, changes in core fucosylation, which emerged in both our glycomic analysis and transcriptomic analysis, was not replicated in the KC mouse model. In addition, several glycomic changes observed in mouse, notably the increase in core 1/3 O-glycans which is observed in other human cancers, are not observed in the human samples (15, 28, 29). Identifying glycans whose expression matches between human patients and the mouse models is a critical step in enabling study of glycans in cancer. This is important in both establishing whether glycans are functionally significant and in setting up pathways for development of new therapeutic agents to target them.

Our analysis pointed to α-2,6-sialylation and its underlying enzyme ST6GAL1 as a potentially important glycan epitope in both systems. Levels of ST6GAL1 protein correlated with severity of human disease and associated with poor prognostic outcomes for pancreatic cancer patients. In line with our sc-RNAseq analysis, staining for ST6GAL1 protein was strongest in epithelial cells, suggesting that ductal or acinar specific expression may significantly contribute to development and/or progression of PDA. Consistent with our observations in patients and the KC mouse model, *in vitro* cell culture models of pancreatic cancer with ST6GAL1 overexpressed promoted chemo-resistance, cell growth and a stem-cell like phenotype (7, 8, 30–32).

Our deletion of ST6GAL1 in only pancreatic specific lineages in the KC mouse enabled assessment of the ductal intrinsic impact of ST6GAL1 in promoting cancer formation and progression. In ST6KC, we observed significant protection against formation of PanIn lesions, an early step in malignant transformation, and reduction of fibrosis, which is associated with immune suppression and resistance to treatment in PDAC patients (33). This is consistent with the function of ST6GAL1 in cell studies (7, 8, 30–32). Overall, histological assessment of our ST6KC mice demonstrates a clear protective role for genetic deletion of ST6GAL1 in PDA.

The protection observed by deletion of epithelial ST6GAL1 suggests that sialyltransferase inhibitors could provide a new therapeutic avenue for PDA. In fact, recent work has shown that pharmacological blockade of α-2,3- and α-2,6-sialyation reduce tumor cell invasion *in vitro* and disease burden in murine models of lung metastasis (34, 35). Our findings in this work suggest that as this emerging class of glycan-targeting inhibitors is developed, the current mouse models will be effective in identifying therapeutic responses to modulating glycans in human pancreatic cancer.

## METHODS

### Animals and Disease Models

C57BL/6 mice were purchased from Jackson Labs (Bar Harbor, ME) and bred in-house. KC mice, which express *Kras^G12D^* in the progenitor cells of the pancreas, were bred in house (36). Both male and female mice were used, and animals were age-matched within each experiment as indicated. All studies were reviewed by the Institutional Animal Care and Use Committee at NYU School of Medicine (IACUC #170513). Experiments were conducted in accordance with the NYU School of Medicine policies on the care, welfare, and treatment of laboratory animals.

### Human Sample Acquisition

All human pancreatic tumor and normal pancreas specimens were collected under an IRB approved protocol (IRB#10-2519) utilizing RedCAP 1253 within the Center for Biospecimen Research and Development at NYU Langone Medical Center. Patient characteristics are given in **Table S1**.

### Murine and Human Sample Processing for Lectin Array

Formalin-fixed paraffin-embedded (FFPE) tissues from mouse and human were prepared for our lectin microarrays, and labeled with AlexaFluor-555 as previously described (15). A pooled reference sample, labeled with AlexaFluor-647, was created for each experiment. Details of sample preparation are given in **Table S2**.

### Lectin Microarray Printing, Hybridization and Analysis

Lectin microarrays were printed as previously described (13). The printlist, including lectin sources, is given in **Table S3**. Equal amounts of sample and reference (5⎧g) were hybridized on each array and data analysis was performed as previously described (37). Additional experimental information can be found in **Table S2**.

### Multiplex Immunofluorescence, Immunohistochemistry, Image Acquisition and Quantification of Murine KC Samples

For histological analysis, murine KC pancreatic tissues were fixed with 10% buffered formalin, dehydrated in ethanol, and embedded with paraffin (FFPE). 5 μm sections of FFPE murine KC were mounted on slides. For immunohistochemical staining, briefly, slides were deparaffinized and stained with hematoxylin and eosin (S3301, Dako, Santa Clara, CA), or Trichrome and Gomore followed by whole-tissue scanning at 40X magnification on an Aperio AT2 (Leica Biosystems, Wetzlar, Germany). Preserved acinar area of KC pancreatic tissues was quantified using PhotoSHOP Software (Adobe Acrobat) and calculated as a fraction of the total pixels of pancreatic tissue. Percent fibrosis was quantified using the total blue positive pixels in Photoshop software (Adobe Acrobat), determined as a fraction of the total pixels of pancreatic tissue.

### Human Tissue Microarray Samples

All staining of human tissues was performed on a Leica Bond RX automated stainer (Leica Microsystems, Inc., Buffalo Grove, IL). To ensure lectin specificity, SNA conjugated to Cy3 (1:300 dilution, Vector Laboratories, cat # CL-1303) was incubated with either 0, 100, 200 mM of beta-D-lactose (Alfa Aesar, cat # H54447) in PBST for 2 hours at 22°C. Liver sections stained with SNA in the absence of beta-D-lactose yielded strong signal. In contrast, SNA staining was blocked by both 100 and 200 mM beta-D-lactose. Prior to duplex staining, Biomax PA2072a TMA underwent deparaffinization and heat retrieval with Bond ER2 buffer (Leica, cat # AR9640). The slide was then incubated with anti-human ST6GAL1 (Proteintech, cat # 14355-1-AP) followed by Opal Polymer HRP Ms + Rb (Akoya Biosciences, cat # ARH1001EA) and tyramide-linked 650 Opal fluorophore (Akoya Biosciences, cat # FP1496001KT) to amplify signal. The antibody complexes were stripped from the tissues with ER2 buffer, leaving the fluorophore covalently attached to ST6GAL1 and the TMA subsequently stained with the SNA-cy3 lectin. Image Acquisition was performed on a Vectra® Polaris multispectral imaging system (Akoya Biosciences, Menlo Park, CA). The fluorophores were spectrally unmixed using InForm software (Akoya Biosciences, Menlo Park, CA) and images exported as tif files for scoring. Cell type identification (epithelial vs stromal) and quantification of number of ST6GAL1 or SNA positive cells per high power field were performed by an independent pathologist and validated by a second scorer for each core.

### Human Tissue RNA Expression Profiling

PDA tumor and adjacent normal tissue mRNA raw expression profiles were downloaded from GEO [accession #: GSE16515 and GSE15471] and normalized in one batch using a GC-content background corrected Robust Multi-array Average (RMA) algorithm (GC-RMA) in *R: A language and environment for statistical computing,* as previously described (38). Hierarchical clustering was performed in GENE-E (https://software.broadinstitute.org/GENE-E/) and adjacent normal tissues clustering with PDA tumors and PDA tumors clustering with adjacent normal were removed. Patients with expression profiles containing the gene of interest in both PDA and adjacent normal groups were utilized for differential expression analysis.

### Cluster Identification and Single Cell Sequencing Analysis

Raw single cell sequencing patient data on was obtained (23). A quality control was then performed on the cells to calculate the number of genes and the proportion of mitochondrial genes for each cell using iCellR R package (v0.99.0) (https://github.com/rezakj/iCellR) and the cells with low number of covered genes (gene-count < 500) and high mitochondrial counts (mt-genes > 0.1) were filtered out. Matrix normalization, geometric library size factor normalization, and gene statistics analysis was performed, as previously published (39). Briefly, genes with high coverage (top 500) and high dispersion (dispersion > 1.5) were chosen and PCA analysis was performed, a second round of PCA was performed based on the top 20 and bottom 20 genes predicted in the first 10 dimensions of PCA to fine tune the results and clustering was performed (iCellR options; clust.method = “kmeans”, dist.method = “euclidean”, index.method = “silhouette”) on principal component with high standard deviation (top 10 PCs) and Uniform Manifold Approximation and Projection (UMAP) was performed on the top 10 PCs. Marker genes for each cluster were determined based on fold change and adjusted p-value (t-test) and average gene expression for each cluster was calculated using iCellR.

### Statistical Analysis and TCGA

Data is presented as mean +/− standard error. Data on gene expression and survival in human tissues was derived from TCGA utilizing cBIOportal (http://cancergenome.nih.gov/). Statistical significance was determined by the Student’s *t* test and the Wilcoxon test using GraphPad Prism 7 (GraphPad Software, La Jolla, CA). p-values <0.05 were considered significant.

## Acknowledgements

We thank members of the Experimental Pathology Research Laboratory, which is partially supported by NIH/NCI 5 P30CA16087 and S10 OD021747 (PerkinElmer/AkoyaBiosciences® Vectra® multispectral imaging system), for their technical support and expertise. E.K. was supported by NIH/NCI grant F30 CA243205. S.C. and L.K.M. were supported in part by funding from DOD-CA171043. Publication of this research was also supported, in part, by funding from the Canada Excellence Research Chairs Program (L. Mahal). We thank Dr. Barbara Bensing (UCSF) for SLBRs and Dr. David Markovitz (University of Michigan) for H84T.

## Author Contributions

(EK)* – KC mouse breeding, data acquisition, data analysis, manuscript preparation (SC)* – lectin microarray analysis, data acquisition, data analysis, manuscript preparation (GB) –data acquisition (EV) – data analysis (CL) –data acquisition, analysis (CH) –data acquisition and analysis (DBS) – manuscript preparation (LKM) – study design, manuscript preparation, data analysis.

