## Supplemental Figure for "Integrated Systems-Analysis of the Human and Murine Pancreatic Cancer Glycomes Reveal a Tumor Promoting Role for ST6GAL1"

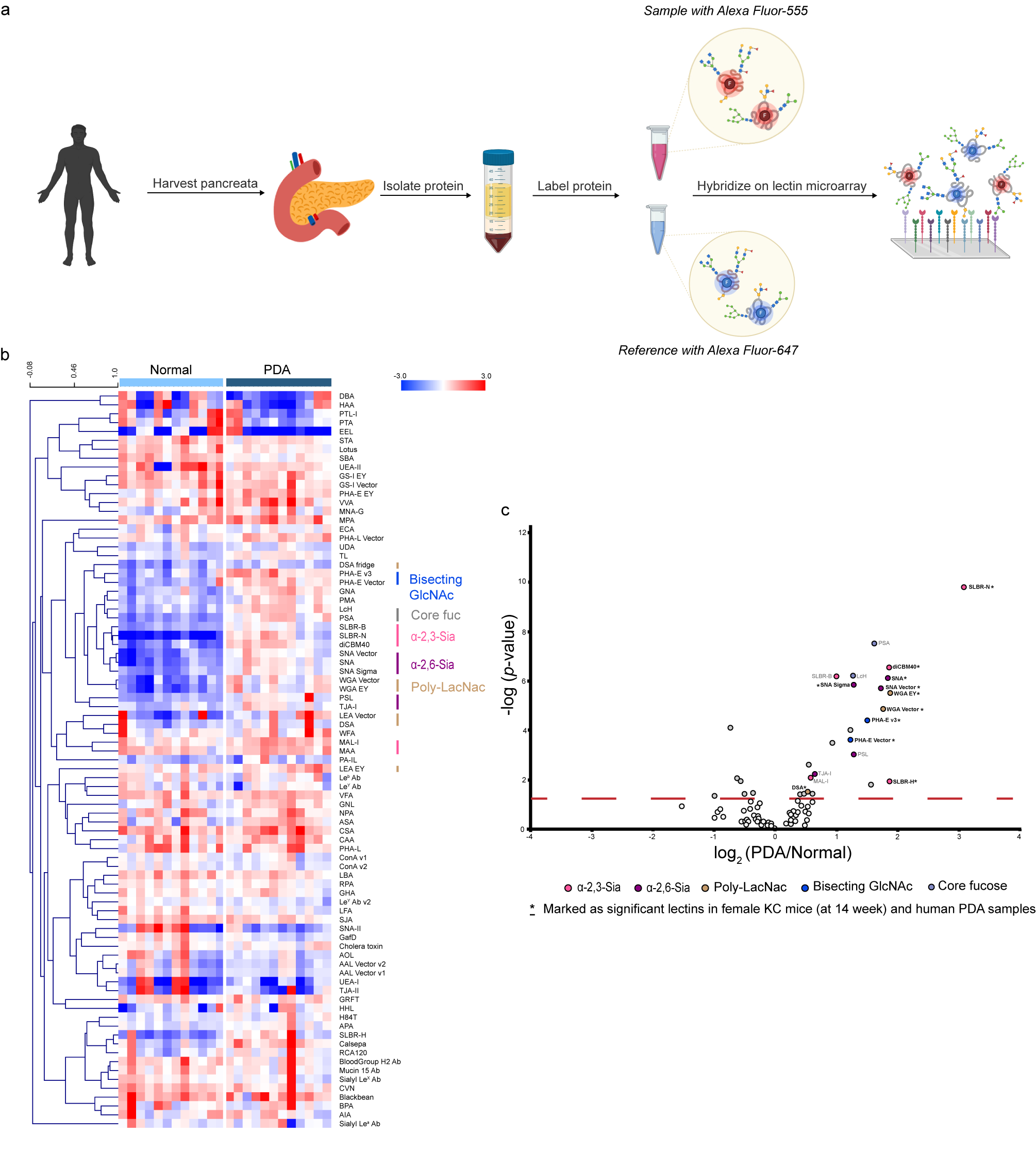

**Supplementary Figure S1. Glycomic analysis of human PDA samples.** a) Workflow of sample preparations for dual-color lectin microarray analysis. Glycoproteins were isolated from pancreata and labeled with Alexa Fluor 555-NHS. A pooled reference was orthogonally labeled with Alexa Fluor 647-NHS. Equal amounts of sample and reference were mixed and hybridized on lectin microarrays (>100 probes). b) Heatmap of human lectin microarray data present with the complete list of lectins. Median normalized log_2_ ratios (Sample (S)/Reference(R)) were ordered by sample type (Normal, n = 12; PDA, n = 12). Red, log_2_(S) > log_2_(R); blue, log_2_(R) > log_2_(S). Lectins binding $\alpha$-2,3-sialosides (pink), $\alpha$-2,6-sialosides (purple), bisecting GlcNAc (navy), poly-*N*-Acetyl-D-Lactosamine (poly-LacNAc, brown), core fucose (charcoal) and Lewis antigens (turquoise) are highlighted to the right of the heatmap. b) Volcano plot analysis showed a decrease in Lewis antigens (turquoise) in human PDA samples compared to normal samples (on the left panel). PDA samples showed an increase in $\alpha$-2,3-sialosides (pink), $\alpha$-2,6-sialosides (purple), bisecting GlcNAc (navy), poly-*N*-Acetyl-D-Lactosamine (poly-LacNAc, brown) and core fucose (slategrey) (on the right panel). Significant lectins (*p*< 0.05) showed in both human PDA samples and female KC mice samples at 14 weeks were bold and marked as asterisk (*). Spot colors correspond to lectin specificity; the dotted line represents a significance cutoff of *p*-value ≤ 0.05 across the biological samples.

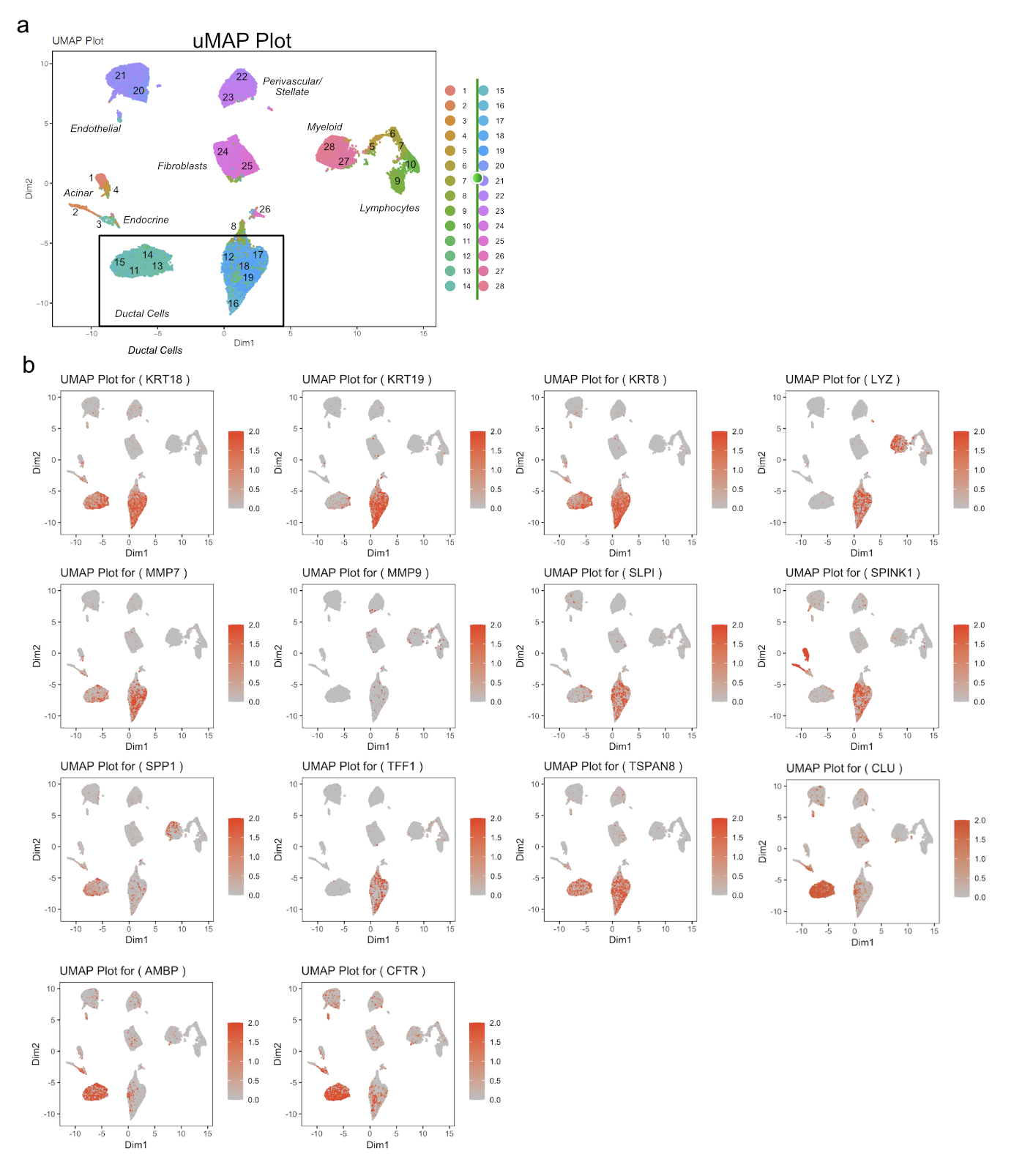
**Supplementary Figure S2. Corroboration of tumor and normal ductal compartments by markers in single cell sequencing.** a) uMAP plot representing all cells isolated from PDAC (n=24) patients and normal pancreata (n=11) pooled on single cell-sequencing and colored and numbered by cluster. b) uMAP plots representing expression of a panel of typical genes that identify cells of ductal origin, including: KRT18, KRT19, KRT8, LYZ, MMP7, MMP9, SLPI, SPINK1, SPP1, TFF1, TSPAN8, CLU, AMBP, and CFTR to validate the identity of ductal clusters in normal and PDAC samples.

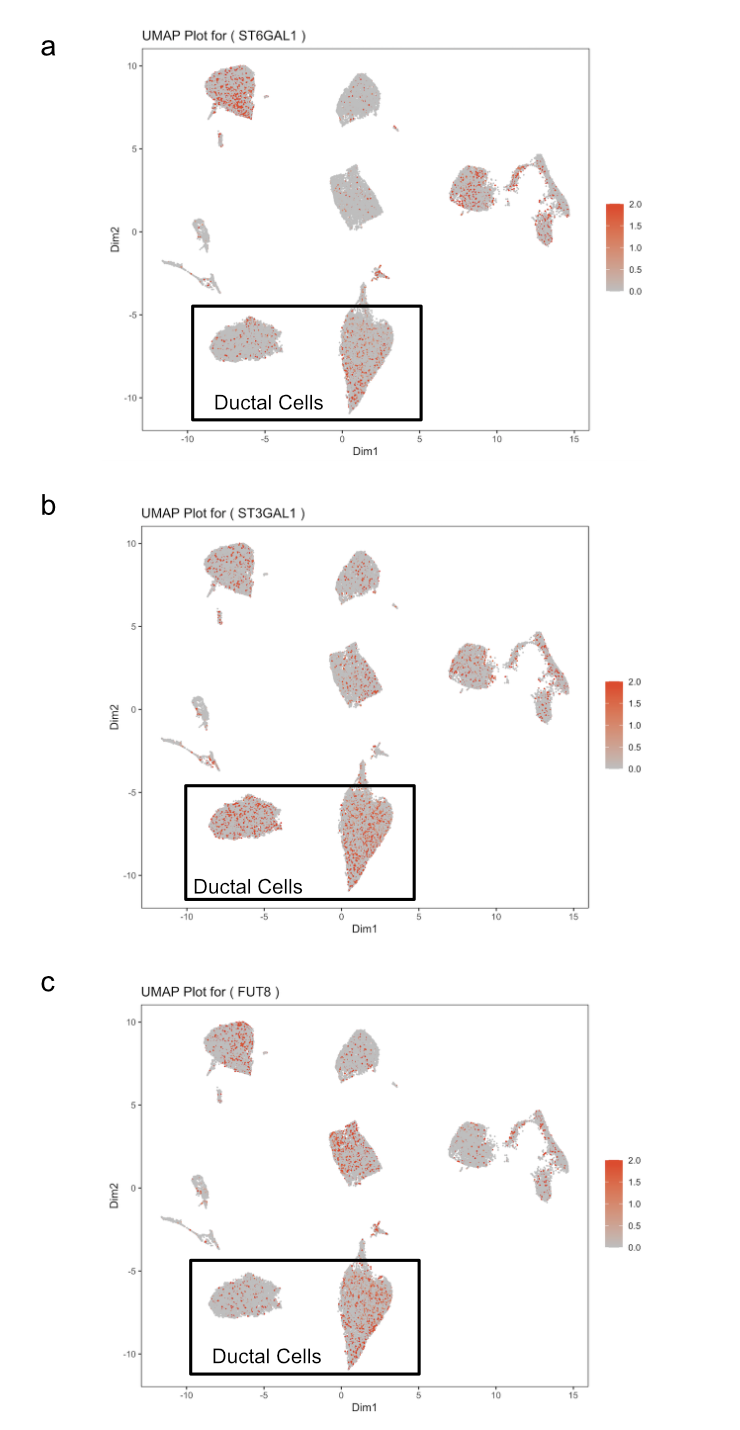

**Supplementary Figure S3. Single cell sequencing analysis of ST6GAL1, ST3GAL1 and FUT8.** a-c) uMAP plots representing cells isolated from PDAC (n=24) patients and normal pancreata (n=11) pooled on single cell-sequencing and colored in red by ST6GAL1, ST3GAL1, and FUT8 mRNA expression. Ductal cell clusters (normal and tumor) are indicated in a box.

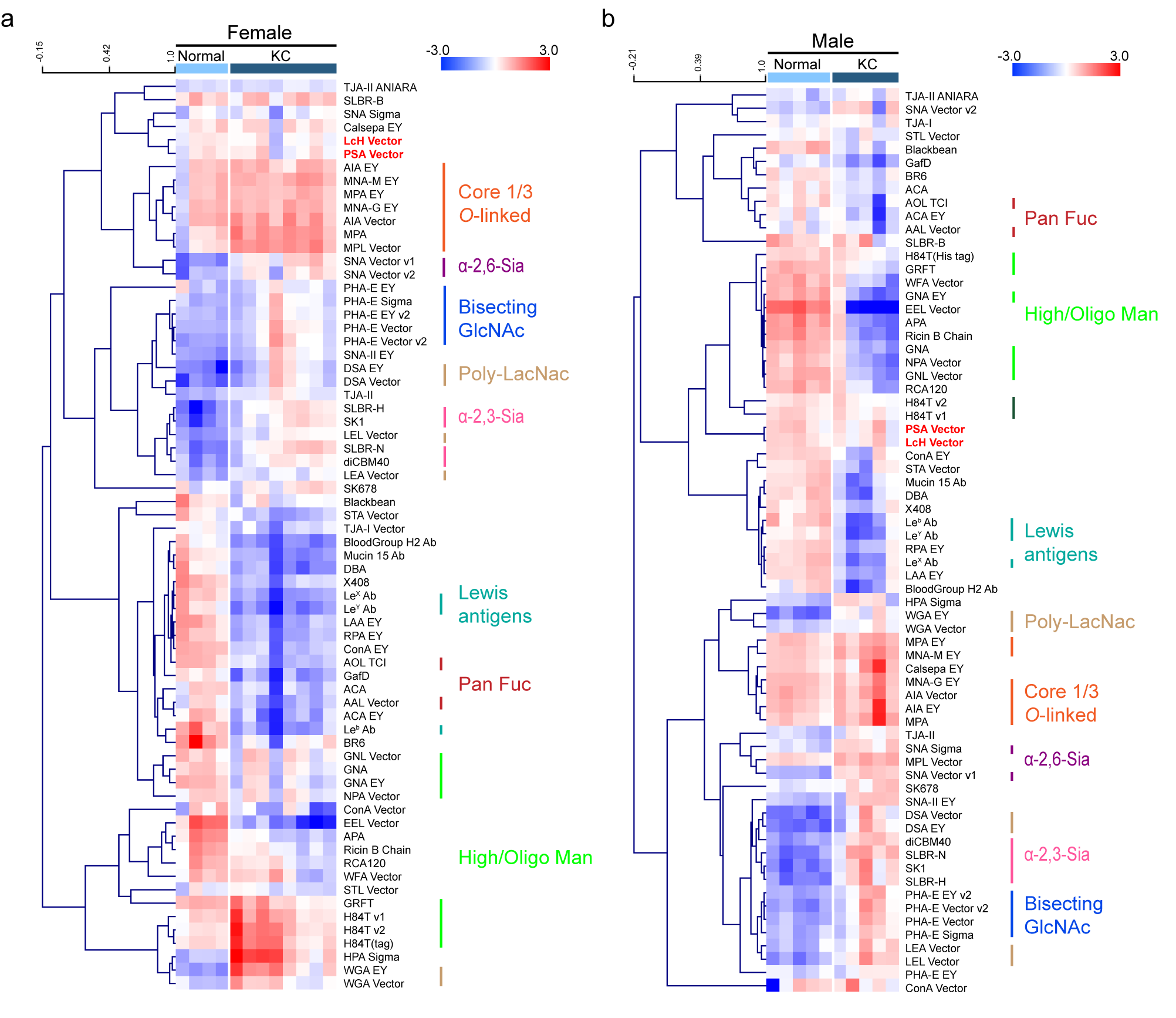
**Supplementary Figure S4. Glycomic analysis of male and female KC mice at 14 weeks of life.** a) Heatmap of female mice lectin microarray data with the complete list of lectins. Median normalized log_2_ ratios (Sample (S)/Reference(R)) were ordered by sample type (Normal, n = 4; KC, n = 8). Red, log_2_(S) > log_2_(R); blue, log_2_(R) > log_2_(S). Lectins binding $\alpha$-2,3-sialosides (pink), $\alpha$-2,6-sialosides (purple), bisecting GlcNAc (navy), poly-*N*-Acetyl-D-Lactosamine (poly-LacNac, brown), high- and oligo-mannose (bright green), core 1/3 *O*-linked glycans (orange), pan fucose (red) and Lewis antigens (turquoise) are highlighted to the right of the heatmap; core fucose binders (LcH, PSA) are bold and colored as red. b) Heatmap of male mice lectin microarray data with the complete list of lectins. Median normalized log_2_ ratios (Sample (S)/Reference(R)) were ordered by sample type (Normal, n = 5; KC, n = 5). Red, log_2_(S) > log_2_(R); blue, log_2_(R) > log_2_(S). Lectins binding $\alpha$-2,3-sialosides (pink), $\alpha$-2,6-sialosides (purple), bisecting GlcNAc (navy), poly-*N*-Acetyl-D-Lactosamine (brown), oligo-mannose (bright green), high-mannose (forest green), core 1/3 *O*-linked glycans (orange), pan fucose (red) and Lewis antigens (turquoise) are highlighted to the right of the heatmap; core fucose binders (LcH, PSA) are bold and colored as red.

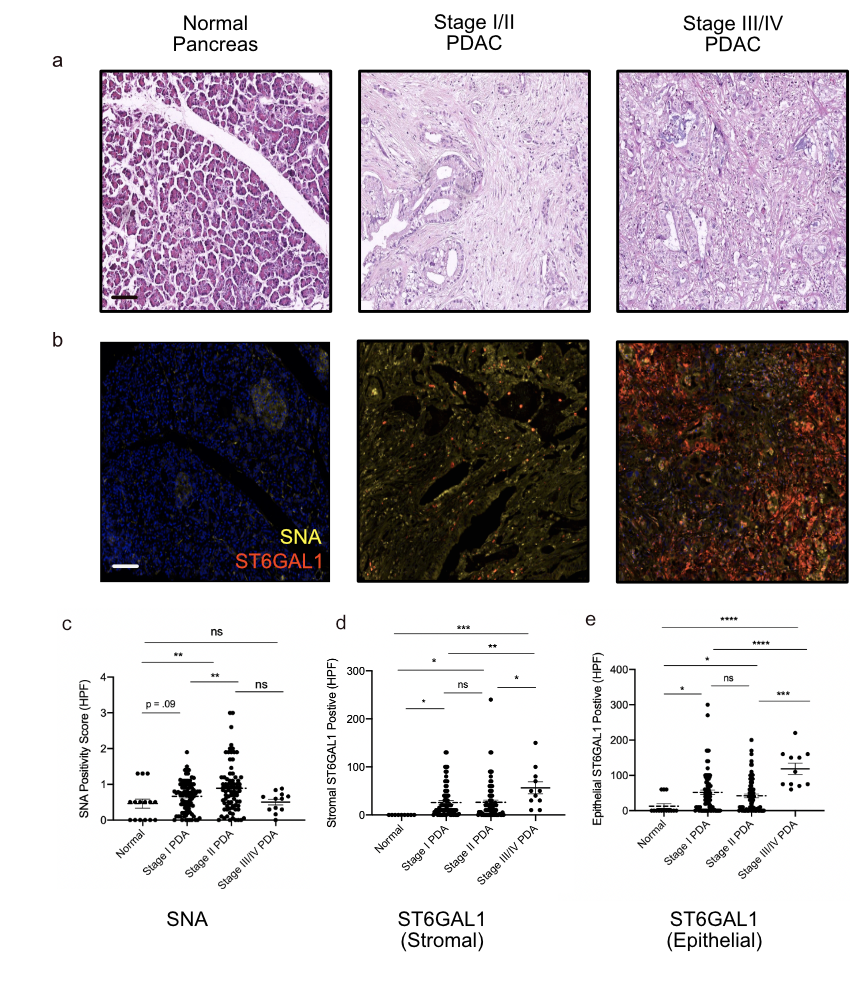

**Supplementary Figure S5. Human TMA Quantification of SNA and ST6GAL1 Cell Type Specific Staining.** a) H&E of normal pancreas (left), stage I pancreatic adenocarcinoma (center), and stage IV PDA (right) stained from a BioMax human tissue microarray. b) Multiplex OPAL IF staining of SNA (yellow), ST6GAL1 (red) and DAPI (blue) on corresponding normal pancreas, Stage I, and Stage IV pancreatic adenocarcinoma from BioMax human tissue microarray. Scale bars represent 100 μm. c) Quantification of SNA positive cells per high powered field based on multiplex IF in normal pancreas compared to tumor samples at each stage of human PDA. d) Quantification of ST6GAL1 positivity in non-epithelial stromal cells in normal pancreas compared to tumor samples at each stage of human PDA e) Quantification of ST6GAL1 positivity in epithelial cells in normal pancreas compared to tumor samples at each stage of human PDA (ns: p > 0.05; *: p < 0.05; **: p < 0.01; ***: p < 0.001; ****: p < 0.0001).
