## Supplemental Table S2 for "Integrated Systems-Analysis of the Human and Murine Pancreatic Cancer Glycomes Reveal a Tumor Promoting Role for ST6GAL1"

**Supplemental** **Table 2. Lectin Microarray Information**

|  | | | **Description** |
| --- | --- | --- | --- |
| 1. **Sample: Glycan-containing sample (e.g. glycan, glycoprotein, cell lysate etc.)** | | | |
| Description of Sample | Glycoproteins extracted from formalin-fixed paraffin-embedded (FFPE) tissues from human PDA patients and KC mouse samples | | |
| Sample preparation protocol | Both human and mouse samples were fixed in 10% neutral buffered formalin. For analysis of tumor samples, hematoxylin and eosin-stained slides were reviewed to ensure the size of tumor. Unstained cut sections were mounted on the slide and macro-dissected to remove containing normal cells. 2 × 20μm sections of each FFPE tissue were scraped into a 1.5mL microcentrifuge tube. 1mL xylene and 200μL 100% ethanol were added to the microcentrifuge tube, incubated for 10 mins at room temperature, centrifuged at 14,000 × g for 3 mins, and supernatant was removed; repeat twice. The deparaffinized tissue were rehydrated with a graded series of ethanol. 1mL 100% ethanol was added to each tissue, incubated for 10 min at room temperature, centrifuged at 14,000 × g for 3 mins, and supernatant was removed. Then 1mL 90% ethanol was added to each tissue, incubated for 10 min at room temperature, centrifuged at 14,000 × g for 3 mins, and supernatant was removed. Next, 1mL 70% ethanol was added to each tissue, incubated for 10 min at room temperature, centrifuged at 14,000 × g for 3 mins, and supernatant was removed. Tissue pellet was heated at 90°C for 30 seconds to remove any remaining ethanol, and allowed to dry at room temperature. The rehydrated tissue was suspended in 200μL 10mM sodium citrate buffer (pH 6.0) and incubated at 95°C for 1 hour. The tissue was centrifuged at 14,000 × g for 5 mins, and supernatant was removed. The tissue was washed with 1× PBS (pH 7.4). The pellet was solubilized with 100μL PBS containing 0.5% Nonidet P-40 (NP40). Samples were sonicated (70% power, 1 min), and incubated on ice for 30 mins; repeat one more time. Sample was centrifuged at 14,000 × g for 15 mins at 4°C, and supernatant was collected for glycomic analysis. | | |
| Labeling protocol for sample detection | Samples are labelled with Alexa Fluor 555-NHS (Thermo Fisher). | | |
| Two-color reference (if used) | A pooled reference samples are labelled with Alexa Fluor 647-NHS (Thermo Fisher). | | |
| Assay protocol | Lectin microarrays are blocked with blocking buffer for one hour at room temperature. Slides are rinsed twice with PBST (0.005%) and once with PBS, then dry the slide using a slide spinner. Each slide was mounted on a 24-well format hybridization cassette (Arrayit), in which each well contains a subarray. To each well, add equal amounts of samples and universal reference, and dilute with PBS and PBST (0.2%) to reach the final volume (150uL). Incubate the slides on an orbital shaker for two hours at room temperature in the dark. After hybridization, wash the arrays with PBST (0.005%) twice for ten minutes, and twice for five minutes. Once finished, remove the slides from the cassette, and immerse the slides in ultrapure water, and dry the slides using a slide spinner. | | |
| **2.** **Lectin Library** | | | |
| General description of the lectin library used in the array | Lectin microarrays are generated in house. | | |
| List of lectins and glycan binding proteins, source, concentration and buffer | Please see **Table S3**. | | |
| Modification of lectins (e.g. biotin) if any. | N/A | | |
| 1. **3.** **Immobilization Surface; e.g., Microarray Slide** | | | |
| Immobilization surface | Nexterion Slide H Barcoded 3D Hydrogel Coated | | |
| Manufacturer | Schott North America | | |
| Custom preparation of surface | N/A | | |
| **4. Array Production** | | | |
| Description of Arrayer | Nano-Plotter 2.1 piezoelectric printer (GeSim, Germany) with cooled microwell plate holder and cooled printing deck | | |
| Lectin deposition | Three replicates of each lectin are printed onto each subarray. | | |
| Printing conditions | Dilute lectins to the pre-determined concentrations in the print buffer (final concentration of print buffer: 0.01% Tween-20, 1mM monosaccharide in PBS; Please see **Table S3** for the concentrations of lectins). Load the mixed solution to the microplate. Before printing, check the humidity of the print chamber. The humidity should be kept around 50% during the entire printing. Ensure both microwell plate holder and printing deck are cooled. Adjust the cooling temperature based on ambient temperature and the temperature of the cooled slide deck surface, preventing moisture building up inside the print chamber. Once printing is complete, allow the slides to dry for at least one hour. | | |
| Array layout | | For each microarray, it contains 24 subarrays (3 columns and 8 rows). In each subarray, triplicates of a lectin are printed, and for a row with five lectins, the spot layout should be 15 columns. The row number depends on how many lectin probes are printed on the arrays (i.e., 110 lectins require 22 rows). | |
| Quality control | | Well-characterized glycoproteins including fetuin, asialofetuin and RNase B are used for quality assurances of the printed microarrays. | |
| 2. **5. Detector and Data Processing** | | | |
| Instrument (scanner, flow cytometer) | Fluorescent Slide Scanner Genepix 4300A (Molecular Devices) | | |
| Instrument settings | Preview the slide to adjust photomultiplier gain (PMT) for each channel (Alexa Fluor-555: 532nm, Alexa Fluor-647: 635nm) so that the signals are not saturated and within the linear detection range. | | |
| Image analysis software | GenePix Pro 7 (Molecular Devices) | | |
| Data processing and statistical analysis | Extracted data is processed for quality checks using Grubbs outlier test with $\alpha$ = 0.05. Log_2_ values of the average signals are median-normalized over the individual subarray in each channel. | | |
| **6.** **Lectin Microarray Data Presentation** | | | |
| Data presentation and interpretation | Hierarchical clustering of the processed data is performed using Pearson Correlation coefficient, and visualized with Multi-experiment Viewer (MeV, v4.8, TM4 Microarray Software Suite). If a lectin’s SNR (signal-to-noise ratio) < 3 for more than one third of the total samples, then this lectin is considered as inactive and excluded from the list. *P*-values are calculated using nonparametric statistical tests, which are generated by R (v3.6.1). | | |
| **7. Data Location** | | | |
| Data Location |  | | |
