## Supplemental Table S3 for "Integrated Systems-Analysis of the Human and Murine Pancreatic Cancer Glycomes Reveal a Tumor Promoting Role for ST6GAL1"

**Supplemental Table 3. Lectins used in microarrays**

| **Lectin** | **Species/Origin** | **Print Conc.**  **(µg/mL)** | **Rough Specificity /Inhibitory monosaccharide** | **Vendor/Source** |
| --- | --- | --- | --- | --- |
| AAL ^a, b^ | *Aleuria aurantia* | 1000 | Fucose | Vector |
| ACA ^a, b^ | *Amaranthus Caudatus* | 1000 | Gal-β1,3-GalNAc | Vector |
| AIA ^a, b^ | *Artocarpus integrifolia* | 500 | β1,3-GalNAc | Vector/EY |
| AMA ^a, b^ | *Allium moly* | 500 | Oligo mannose | EY |
| Anti-B.G.H2 ^a, b^ | MAb mouse IgM [A46-B/B10] | undiluted | Blood group H2 antigen | Santa Cruz Biotechnology |
| Anti-Forssman ^a^ | MAb Rat IgM [117C9] | undiluted | Forssman Antigen | Abcam |
| Anti-Lewis B ^a, b^ | IgM [T218] | undiluted | Lewis B | Sigma |
| Anti-Lewis X ^a, b^ | MAb mouse IgM [P12] | undiluted | Lewis X | Abcam |
| Anti-Lewis Y ^a, b^ | MAb mouse IgM [F3] | undiluted | Lewis Y | Abcam |
| Anti-MUC5AC human ^a, b^ | Mab mouse IgG1 [CLH2] | undiluted | human MUC5AC | Sigma |
| Anti-MUC5AC mouse ^a^ | Goat polyclonal to mouse MUC5AC | undiluted | mouse MUC5AC | LSBio |
| Anti-Mucin 15 ^a, b^ | Mab mouse IgG1 [H-5] | undiluted | Mucin 15 | Santa Cruz Biotechnology |
| Anti-Sialyl Lewis A ^a^ | Mab mouse IgG1 | undiluted | Sialyl Lewis A | Abcam |
| Anti-Sialyl Lewis X ^a^ | Mab mouse IgM | undiluted | Sialyl Lewis X | Abcam |
| AOL ^a, b^ | *Aspergillus oryzae* | 1000 | Fucose | TCI America |
| APA ^a, b^ | *Abrus precatorius* | 500 | Gal-β1,3-GalNAc / Lac | EY |
| ASA ^a, b^ | *Allium sativum* | 1000 | Mannose | EY |
| Blackbean ^a, b^ | *Blackbean crude* | 1000 | GalNAc | EY |
| BPA ^a, b^ | *Bauhinia purpurea* | 500 | β-Gal / β-GalNAc | Vector |
| BR6 ^b^ | unknown (from unpublished work) | 480 | under investigation | Gift from Dr. Barbara Bensing |
| CA ^b^ | *Colchicum autumnale* | 1200 | Bi-antennary N-linked glycans | EY |
| CAA ^a^ | *Caragana arborescens* | 1000 | Bi-antennary N-linked glycans | EY |
| Calsepa ^a, b^ | *Calystegia sepium* | 1000 | Bisecting N-linked glycans | EY |
| CCA ^a^ | *Cancer antennarius* | 1000 | 9-O-Acetly sialylation / 4-O-Acetyl sialylation | EY |
| Cholera Toxin ^a, b^ | *Vibrio cholerae* | 1000 | GM1 ganglioside | Sigma |
| Con A ^a, b^ | *Canavalia ensiformis* | 1000 | Tri-mannose core | EY/Vector |
| CSA ^a, b^ | *Cystisus scoparius* | 1000 | Terminal GalNAc | EY |
| DBA ^a, b^ | *Dolichos Biflorus* | 1000 | GalNAc | Vector |
| diCBM40 ^a, b^ | engineered NanI from *Clostridium perfringens* | 1000 | α Sialylation | Generated in house |
| DSA ^a, b^ | *Datura stramonium* | 500 | LacNAc | EY/Vector |
| ECA ^a, b^ | *Erythrina cristagalli* | 1000 | LacNAc | Vector |
| EEL/EEA ^a, b^ | *Eunonymus europaeus* | 1000 | Blood Group B | Vector/EY |
| GafD ^a, b^ | recombinant GafD from *Escherichia coli* | 1000 | GlcNAc | Generated in house |
| GHA ^a^ | *Glechoma hederacea* | 500 | GalNac | EY |
| GNA/GNL ^a, b^ | *Galanthus nivalis* | 1500 | Oligo mannose | Vector/EY |
| GS-I ^a, b^ | *Griffonia simplicifoia-I* | 1000 | α-Gal / Lac | Vector/EY |
| GS-II ^a, b^ | *Griffonia simplicifoia-II* | 1000 | GlcNAc | Vector |
| GS-IB4 ^a, b^ | *Griffonia simplicifoia-I, isolectin B4* | 2000 | Gal | Vector |
| H84T ^a, b^ | *Banana lectin* | 1000 | High mannose | Gift from Dr. David Markovitz |
| HAA ^a, b^ | *Homarus americanus* | 1000 | Terminal GalNAc | EY |
| HHL ^a, b^ | *Hippeastrum Hybrid* | 1500 | Oligo/High mannose | Vector |
| HPA ^a, b^ | *Helix pomatia* | 1000 | Blood Group A | Sigma/EY |
| LAA ^b^ | *Laburnum alpinum* | 900 | GlcNAc | EY |
| LBA ^a^ | *Phaseolus lunatus* | 1000 | Blood Group A | EY |
| LcH ^a, b^ | *Lens Culinaris* | 1000 | Core Fucose | Vector |
| LEA/LEL ^a, b^ | *Lycopersicon esculentum* | 1000 | GlcNAc | Vector/EY |
| LFA ^a^ | *Limax flavus* | 500 | α Sialylation | EY |
| Lotus ^a, b^ | *Lotus tetragonolobus* | 1000 | Fucose | Vector |
| MAA ^a^ | *Maackia amurensis* | 500 | Sialylation/Sulfation | EY |
| MAL-I ^a, b^ | *Maackia amurensis-I* | 2000 | Sialylation/Sulfation | Vector |
| MAL-II ^a, b^ | *Maackia amurensis-II* | 2000 | Sialylation/Sulfation | Vector |
| MNA-G ^a, b^ | *Morus nigra Morniga G* | 1000 | GalNAc | EY |
| MNA-M ^b^ | *Morus nigra Morniga M* | 1000 | Oligo mannose / Gal | EY |
| MPA/MPL ^a, b^ | *Maclura pomifera* | 1000 | β1,3-GalNAc | Vector |
| NPA ^a, b^ | *Narcissus pseudonarcissus* | 1000 | Oligo mannose | Vector |
| PA-I ^b^ | *Pseudomonas aeruginosa* | 1000 | Gal | Sigma |
| PA-IL ^a^ | *bacteria* | 1000 | GalNAc | Generated in house |
| PHA-E ^a, b^ | *Phaseolus vulgaris Erythroagglutinin* | 1000 | Bisecting GlcNAc | Vector/EY/Sigma |
| PHA-L ^a, b^ | *Phaseolus vulgaris Leukoagglutinin* | 1000 | β1,6 Branching N-Link glycans | Vector/EY/Roche |
| PMA ^a^ | *Polygonatum multiflorum* | 500 | Oligo mannose | EY |
| PNA ^a, b^ | *Arachis hyogaea* | 1000 | Gal-β1,3-GalNAc | Vector/EY |
| PSA ^a, b^ | *Pisum sativum* | 1000 | Core Fucose | Vector |
| PSL ^a^ | *Polyporus squamosus* | 1000 | α2,6 sialylation | EY |
| PTA ^a, b^ | *Psophocarpus tetragonolobus* | 500 | Blood Groups | EY |
| PTL-I ^a, b^ | *Psophocarpus tetragonolobus-I* | 1500 | Blood Group A | Vector |
| PTL-II ^a, b^ | *Psophocarpus tetragonolobus-II* | 1000 | α2 Fucose | Vector |
| RCA120 ^a, b^ | *Ricinus Communis Agglutinin I* | 1000 | Gal / Lac | Vector |
| rCVN ^a^ | *recombinant Cyanovirin* | 1000 | High mannose | Gift from Dr. Barry O'Keefe |
| rGRFT ^a, b^ | *recombinant Griffithsin* | 1000 | High mannose | Gift from Dr. Barry O'Keefe |
| Ricin B Chain ^a, b^ | *Ricinus communis* | 1000 | Gal | Vector |
| RPA ^a, b^ | *Robinia pseudoacacia* | 500 | Complex N-link glycans | EY |
| rSVN ^a^ | *recombinant Scytovirin* | 1000 | High mannose | Gift from Dr. Barry O'Keefe |
| SBA ^a, b^ | *Glycine max* | 1000 | LacdiNAc | Vector |
| SJA ^a, b^ | *Sophora japonica* | 1000 | LacdiNAc | Vector |
| SK1 ^b^ | *Streptococcus sanguinis SK1* | 1800 | α2,3 sialylation | Gift from Dr. Barbara Bensing |
| SK678 ^b^ | *Streptococcus sanguinis SK678* | 450 | α2,3 sialylation | Gift from Dr. Barbara Bensing |
| SLBR-B ^a, b^ | *Streptococcus gordonii M99* | 1000 | α2,3 sialylation | Gift from Dr. Barbara Bensing |
| SLBR-H ^a, b^ | *Streptococcus gordonii DL1* | 2000 | α2,3 sialylation | Gift from Dr. Barbara Bensing |
| SLBR-N ^a, b^ | *Streptococcus gordonii UB10712* | 1000 | α2,3 sialylation | Gift from Dr. Barbara Bensing |
| SNA ^a, b^ | *Sambucus nigra* | 500/1000 | α2,6 sialylation | Vector/Sigma |
| SNA-II ^a, b^ | *Sambucus nigra-II* | 1000 | α2 Fucose /oligo mannose | EY |
| STA/STL ^a, b^ | *Solanus tuberosum* | 500 | GlcNAc | Vector |
| TJA-I ^a, b^ | *Trichosanthes japonica-I* | 1000 | α2,6 sialylation | TCI |
| TJA-II ^a, b^ | *Trichosanthes japonica-II* | 1000 | α2 Fucose | NorthStar Bioproducts/Aniara Diagnostica |
| TL ^a, b^ | *Tulipa sp.* | 700 | GlcNAc | EY |
| UDA ^a^ | *Urtica dioica* | 1000 | GlcNAc / Oligo mannose | EY |
| UEA-I ^a, b^ | *Ulex europaaeus-I* | 1000 | α2 Fucose | Vector |
| UEA-II ^a, b^ | *Ulex europaaeus-II* | 2000 | GlcNAc | Vector |
| VFA ^a, b^ | *Vicia faba* | 1000 | GlcNAc | EY |
| VVA ^a, b^ | *Vicia villosa* | 1000 | Terminal GalNAc | Vector/EY |
| VVA(man) ^a, b^ | *Vicia villosa* | 500 | Mannose | Vector/EY |
| X408 ^b^ | unknown (from unpublished work) | 1000 | under investigation | Gift from Dr. Barbara Bensing |
| WFA ^a, b^ | *Wisteria floribunda* | 1000 | GalNAc-β1,4 | Vector |
| WGA ^a, b^ | *Triticum vulgare* | 1000 | GlcNAc | Vector/EY |

^a^ : lectins printed in human lectin microarrays

^b^ : lectins printed in mouse lectin microarrays
